# Single Nuclei RNA Sequencing of the Gastrocnemius Muscle in Peripheral Artery Disease

**DOI:** 10.1101/2023.06.05.543809

**Authors:** Caroline Pass, Victoria Palzkill, Jianna Tan, Kyoungrae Kim, Trace Thome, Qingping Yang, Brian Fazzone, Scott T. Robinson, Kerri A. O’Malley, Feng Yue, Salvatore T. Scali, Scott A. Berceli, Terence E. Ryan

**Author notes:** Correspondence: Terence E. Ryan, PhD: 1864 Stadium Rd, Gainesville, FL, 32611.; Twitter: @TerenceRyan_PhD.

## Abstract

**Background:** Lower extremity peripheral artery disease (PAD) is a growing epidemic with limited effective treatment options. Herein, we provide a single nuclei atlas of PAD limb muscle to facilitate a better understanding of the composition of cells and transcriptional differences that comprise the diseased limb muscle.

**Methods:** We obtained gastrocnemius muscle specimens from 20 PAD patients and 12 non-PAD controls. Nuclei were isolated and single nuclei RNA sequencing (snRNAseq) was performed. The composition of nuclei was characterized by iterative clustering via principal component analysis, differential expression analysis, and the use of known marker genes. Bioinformatics analysis was performed to determine differences in gene expression between PAD and non-PAD nuclei, as well as subsequent analysis of intercellular signaling networks. Additional histological analyses of muscle specimens accompany the snRNAseq atlas.

**Results:** snRNAseq analysis indicated a fiber type shim with PAD patients having fewer Type I (slow/oxidative) and more Type II (fast/glycolytic) myonuclei compared to non-PAD, which was confirmed using immunostaining of muscle specimens. Myonuclei from PAD displayed global upregulation of genes involved in stress response, autophagy, hypoxia, and atrophy. Subclustering of myonuclei also identified populations that were unique to PAD muscle characterized by metabolic dysregulation. PAD muscles also displayed unique transcriptional profiles and increased diversity of transcriptomes in muscle stem cells, regenerating myonuclei, and fibro-adipogenic progenitor (FAPs) cells. Analysis of intercellular communication networks revealed FAPs as a major signaling hub in PAD muscle, as well as deficiencies in angiogenic and bone morphogenetic protein signaling which may contribute to poor limb function in PAD.

**Conclusions:** This reference snRNAseq atlas provides a comprehensive analysis of the cell composition, transcriptional signature, and intercellular communication pathways that are altered in the PAD condition.

## INTRODUCTION

Peripheral artery disease (PAD) is a worldwide epidemic that estimated to impact more than 220 million people ^1, 2^ and is expected to continue increasing in prevalence over the coming decades. Lower extremity PAD is caused by atherosclerosis in the peripheral vessels and results in decreased blood flow to the legs. Risk factors for PAD include smoking, diabetes, dyslipidemia, hypertension, and chronic kidney disease. Patients with lower extremity PAD commonly experience a progressive deterioration of physical function and walking performance that reduce quality of life^3^ and have been associated with mortality risk^4–6^. While evidenced-based guidelines for the management of PAD recommend several medications that reduce cardiovascular events (antiplatelets, statins, antihypertensives), Cilostazol, which was approved for use in the United States in 1999, is the only medication available to attenuate the limb symptoms^7^. Recently, supervised exercise training has been established as an effective therapy that improves walking performance in PAD patients^8–10^, although participation in such programs is not widespread in the PAD patient population^11^. To date, experimental protein, gene, and cell-based therapies aimed to improve blood flow or angiogenesis have failed to improve outcomes in clinical trials^12–18^. For patients with more severe PAD symptomology that limits quality of life, endovascular and open revascularization surgical approaches are commonly employed, but unfortunately the rate of adverse limb events remains high^19^.

Skeletal muscle pathology in PAD patients has garnered significant attention^20, 21^ and evidence has indicated that muscle size and function have strong associations with patient outcomes including quality of life, walking performance, and mobility decline^22–30^. The emergence of single cell and single nuclei technologies has enabled detailed characterization of cell types and phenotypic diversity in complex tissues including skeletal muscle^31, 32^. These technologies have also facilitated a deeper understanding of the intercellular signaling networks that may drive both pathological and beneficial adaptations within muscle^33, 34^. Considering the complex network of cell types and pathological microenvironment within the PAD limb, the development of a reference atlas of the PAD limb muscle at a single cell resolution has the potential to elucidate novel mechanisms of disease pathobiology as well as new therapeutic targets that may have better efficacy in improving limb function.

Here, we applied single nuclei RNA sequencing to 57,364 high quality nuclei obtained from the gastrocnemius muscles of 20 PAD patients and 12 non-PAD controls to elucidate the cell-specific transcriptional signatures of the PAD limb. These data reveal phenotypic diversity within several cell populations including myonuclei, muscle stem cells (MuSCs, regenerating myonuclei, and fibro-adipogenic progenitor (FAPs) cells that were unique to the PAD patient. We also uncovered aberrant intercellular communication, including deficient bone morphogenetic protein (BMP) signaling, that likely contribute to the atrophic condition observed in the gastrocnemius muscle of PAD patients. This single nucleus atlas provides a foundation for understanding the complex pathological environment within the PAD limb muscle and intercellular communications that could help explain the lack of effective therapeutic advancement to improve mobility impairment in PAD.

## MATERIALS AND METHODS

### Human ParGcipants

Gastrocnemius muscle specimens were collected from non-PAD adults, PAD patients via percutaneous muscle biopsy using sterile procedures as previously described^28, 35^. A portion of the muscle was quickly trimmed of fat/connective tissue and snap frozen in liquid nitrogen for single nuclei RNA sequencing analysis. Another portion of muscle was frozen in liquid nitrogen cooled isopentane for histological analysis. This study was approved by the institutional review boards at the University of Florida and the Malcom Randall VA Medical Center (Gainesville, FL). All study procedures were carried out according to the Declaration of Helsinki and participants were fully informed about the research and informed consent was obtained.

### Skeletal muscle histology

Skeletal muscle histopathology was assessed using Masson’s trichrome staining. Frozen transverse sections (8µm) of the muscle specimen were cut and mounted on microscope slides. Slides were fixed using 4% PFA for one hour at room temperature and stored in Bouin’s Solution overnight at RT. Slides were removed from Bouin’s solution (Cat No. HT15, Sigma-Aldrich) and placed in a running lukewarm tap water bath for two minutes, immediately following slides were continuously dipped in distilled water for one minute. Slides were then placed in equal parts Weigert’s Iron Hematoxylin Solution A: Solution B (Cat No. HT1079, Sigma-

Aldrich) for three minutes and then placed in running warm tap water bath for 15 minutes. Slides were then continuously dipped in distilled water for one minute and then placed in Biebrich Scarlet Acid Fuchsin (Cat No. HT15, Sigma-Aldrich) solution for one minute. Immediately following, slides were continuously dipped in 3×1 minute fresh distilled water baths. Slides were then placed in one-part Phosphotungstic Acid, one-part Phosphomolybdic Acid (Cat No. HT15, Sigma-Aldrich) and two parts distilled water for ten minutes. Amer ten minutes, slides were removed, and excess liquid was removed by gently tapping slides on a kim wipe and immediately placed in Aniline Blue (Cat No. HT15, Sigma-Aldrich) solution for 15 minutes. Slides were then continuously dipped in 3×1 minute in fresh distilled water baths. Slides were placed in 1% Glacial Acetic Acid for two minutes and then rinsed in 2×1 minute fresh distilled water baths. Slides were dehydrated for three minutes each in increasing concentrations of ethanol (70%-100%). Following ethanol washed slides were dipped five times in a 1:1 xylene:ethanol solution and then placed in xylene for ten minutes. Cover slips were mounted using toluene. Slides were imaged at 20× magnification with an Evos FL2 Auto microscope (ThermoFisher Scientific). Fibrosis was quantified as a percentage of total muscle area.

### Immunofluorescence microscopy

Skeletal muscle fiber cross-sectional area (CSA) and myosin heavy chain isoform were assessed using immunofluorescence microscopy. Frozen transverse sections (8µm) of the muscle specimen were cut and mounted on microscope slides. To label the sarcolemma, the sections were incubated with primary antibody against to laminin (Developmental Studies Hybridoma Bank, University of Iowa, Cat. No. 2E8, 1:100 dilution). Additionally, sections were labeled with primary antibodies against myosin heavy chain (MyHC) isoforms as follows: MyHC I (Cat. No. BA-D5, 1:100 dilution), MyHC IIa (Cat. No. SC-71, 1:500 dilution), and MyHC IIx (Cat. No. 6H1, 1:100 dilution) (all from the Developmental Studies Hybridoma Bank, University of Iowa) overnight at 4°C. Next, slides were washed by 4 × 5-minutes with phosphate buffered saline (PBS) and subsequently incubated with appropriate Alexa-Fluor-conjugated secondary antibodies (all 1:300 dilution) for one hour at room temperature. Following 4 × 5-minute washes with 1× PBS, coverslips were mounted with Vectashield hardmount (Vector Laboratories, Cat. No. H-1400). Slides were imaged at ×20 magnification with an Evos FL2 Auto microscope (ThermoFisher Scientific), and tiled images of the entire muscles were obtained for analysis. The proportion of myofiber types (myosin heavy chain) were analyzed by manually counting fiber numbers in ImageJ by a blinded investigator. Quantification of myofiber cross-sectional area (CSA) was performed using Myosight, an automated macro developed in Fiji/ImageJ^36^.

To assess vascular density in muscle specimens, sections were incubated with primary antibodies to label capillaries (anti-CD31, Abcam, Cat. No. ab7388, 1:200 dilution) and arterioles (anO-smtioth muscle actin, ThermoFisher Scientific, Cat. No. 14-9760-82, 1:400 dilution) overnight at 4°C. Next, slides were washed by 4 × 5-minutes with phosphate buffered saline (PBS) and subsequently incubated with appropriate Alexa-Fluor-conjugated secondary antibodies (all 1:300 dilution) for one hour at room temperature. Following 4 × 5-minute washes with 1× PBS, coverslips were mounted with Vectashield hardmount (Vector Laboratories, Cat. No. H-1400). Slides were imaged at ×20 magnification with an Evos FL2 Auto microscope (ThermoFisher Scientific), and tiled images of the entire muscles were obtained for analysis. To quantify the density of capillaries and arterioles, the tiled images were thresholded and counted by a blinded investigator and the vessel density was normalized to the area of the muscle section.

### snRNA sequencing

Nuclei were isolated from the gastrocnemius muscle specimens of patients with and without PAD using the Chromium nuclei isolation kit (10x Genomics, Cat. No. PN-100494). snRNAseq libraries were generated using Chromium Next GEM Single Cell 3’ HT Reagent kits v3.1 (10x Genomics, Cat. No. PN-100370). Pooled libraries were sequenced on an Illumina NovaSeq 6000 with a S4 flow cell and 2 x 150bp pair-end reads (Azenta Life Sciences).

### snRNA-seq data processing and analysis

Datasets were processed through 10x Genomics pipeline, CellRanger count (v7.0.0). Alignment was performed to the Human (GRCh38) 2020-A reference genome. Ambient RNA signal were removed for each sample using the default SoupX (v.1.6.0)^37^ workflow (https://github.com/constantAmateur/SoupX). Samples were pre-processed independently using Seurat (v4.1)^38^ workflow (https://github.com/saOjalab/seurat). Nuclei with greater than 4,000 and less than 400 features were removed from all datasets. Nuclei with greater than 5% of unique molecular identifiers derived from mitochondrial genes were removed. Amer pre-processing, DoubletFinder (v2.0.3)^39^ was used to identify and remove putative doublets in each dataset. Pre-processed and filtered Seurat objects were integrated using the Seurat function FindIntegrationAnchors and the anchors were run through IntegrateData to generate a new Seurat object. The integrated object was then scaled, and the principal component analysis was performed, including up to 40 principal components. A shared nearest neighbor (SNN) graph was built using the first 40 principal components through Seurat’s FindNeighbors. Cells were clustered with FindClusters with a resolution of 0.8. A dimensional reduction UMAP was generated through the Seurat function RunUMAP for visualization. FindAllMarkers identified top marker genes for each cluster. Cluster identioes were annotated based on top marker genes ad expression of known marker genes based on published literature. To determine differentially expressed genes (DEG) between samples in each cluster, a metadata column was first generated to characterize nuclei based on cluster identity and sample type. Object idents were then modified to include both cluster identity and sample. Finally, FindMarkers was run to compare DEGs between samples based on cluster identity. DEG lists were generated with up and down regulated genes for each cluster identity using a Wilcoxon Rank Sum test and only considering genes with > log2(0.25) fold-change and expressed in at least 25% of cells in the cluster. *P*-values were corrected for false-discovery (FDR) and reported as adjusted *P*-values. Gene set enrichment analyses were performed through ClusterProfiler^40^ function gseGO, using DEG lists. Gene sets enriched in PAD were selected based on *q* value and normalized enrichment score (NES). Myonuclei populations were subset and re-clustered independently following Seurat standard workflow. SNN graphs were built using the top eleven principal components and clustering with a resolution of 0.5. MuSC, Regenerating myonuclei, vascular nuclei, and FAPs were subset out for PAD and non-PAD and analysis was performed in Scanpy^41^ (v1.9.2) using the Leiden algorithm to re-cluster non-PAD and PAD nuclei populations independently. scFates^42^ (v1.0.2) analysis was used to perform trajectory analysis on MuSCs and Regenerating myonuclei. In this analysis, the gene *PAX7* was used as the root for trajectory analysis and genes with significant changes (adjusted *P*-value less than 0.05) as a function of pseudtiome were determined. Normalized cell count data and cell group information were obtained from Seurat non-PAD and PAD objects to generate independent

CellChat^43^ (v1.6.1) objects. The CellChatDB ligand-receptor database was used for the cell-cell communication analysis. Downstream cell-cell communication analyses were performed independently following the CellChat workflow on non-PAD and PAD objects. Non-PAD and PAD CellChat objects were merged for interaction comparisons. All sequencing data have been deposited in the Gene Expression Omnibus (https://www.ncbi.nlm.nih.gov/geo/) under accession number GSE233882.

### StaGsGcal analysis

All data are presented as violin plots with the median and interquartile ranges. When possible, individual data points are shown within the violin plots. Normality of data was tested with the Shapiro-Wilk test and inspection of QQ plots. Data involving comparisons of normally distributed data from two groups were analyzed using an unpaired student’s *t*-test. Data involving comparisons of non-normally distributed data from two groups were analyzed using the Kolmogorov-Smirnov test. For snRNAseq, false discovery rate corrected *P-*value were used to determine genes that were statistically different between non-PAD and PAD nuclei populations. In all cases, *P* < 0.05 was considered statistically significant. All statistical testing was conducted using R-studio, Python, or GraphPad Prism somware (version 9.0).

## RESULTS

### Single nuclei RNA sequencing (snRNAseq) reveals differences in cell populaGons and transcripGonal profiles in the gastrocnemius of PAD paGents

To compare different nuclei populations in PAD and non-PAD limb muscles, we performed snRNAseq on nuclei isolated from the gastrocnemius muscle of 20 PAD patients (7 female) and 12 non-PAD (5 female) controls (**Figure 1A**). Non-PAD participants were of similar age, BMI, and had similar medication usage as PAD patients (**Table 1**). PAD patients had greater prevalence of strong risk factors for the disease including hypertension (*P*=0.084), hyperlipidemia (*P*=0.030), and coronary artery disease (*P*=0.005) when compared to non-PAD participants. Ankle brachial index was significantly lower in PAD patients (0.62 ± 0.15 vs. 1.08 ± 0.09 in non-PAD, *P*<0.001). From non-PAD muscles, we captured 21,417 nuclei, and from PAD patient muscle we captured 35,947 nuclei with high integrity (**Supplemental Figure 1**). A median of 2,391 genes per nuclei were sequenced from non-PAD muscle, whereas a median of 2,057 genes per nuclei were sequenced from PAD muscle. The total number of genes detected was 31,671 and 31,010 in non-PAD and PAD muscle respectively.

**Figure 1.**
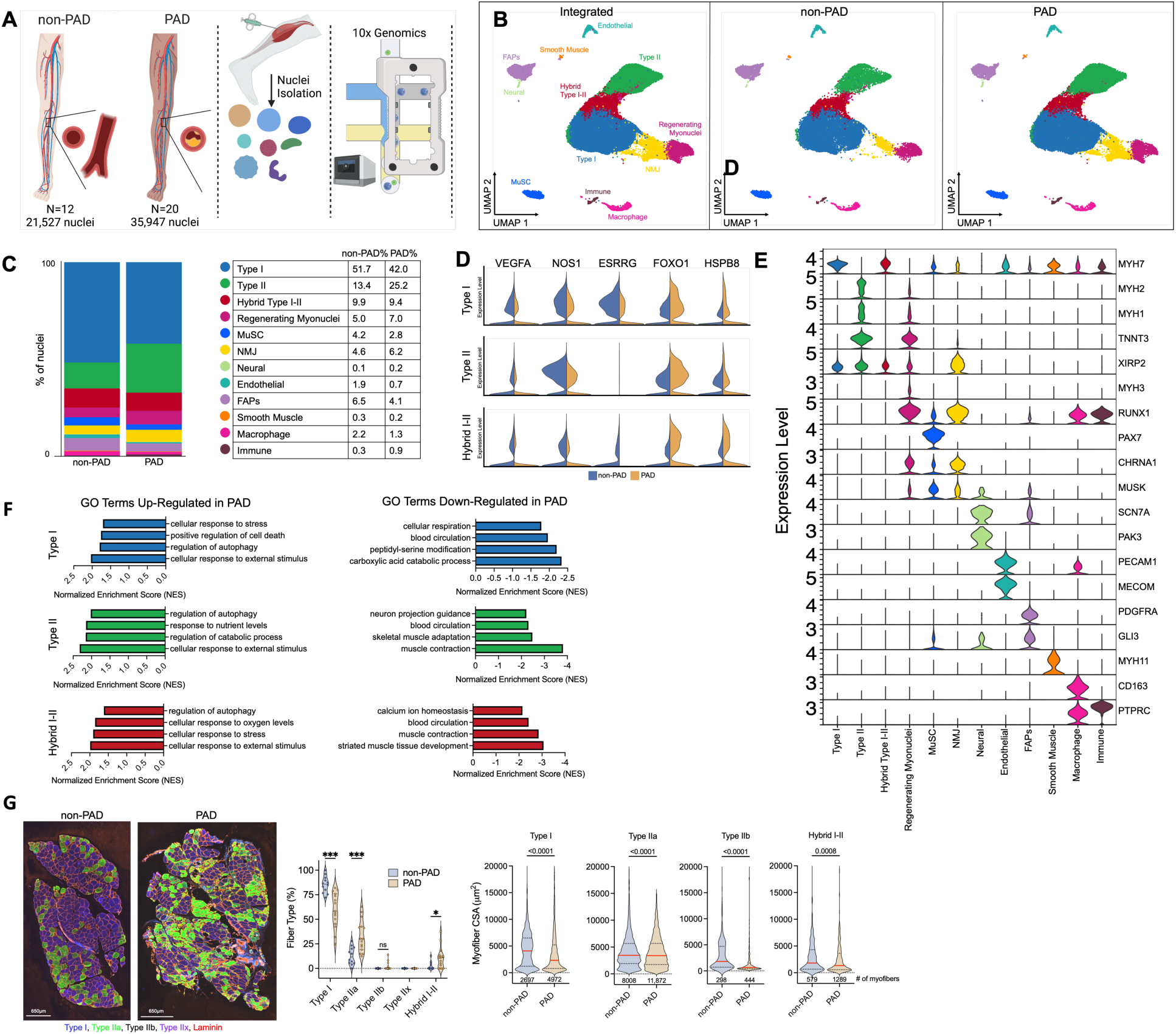
Single-nuclei transcripGonal profiling of the gastrocnemius muscle in non-PAD controls and PAD paGents. (**A**) Schematic of snRNAseq experiment. (**B**) UMAP visualization of integrated datasets to identify clusters and UMAPs for non-PAD and PAD conditions. (**C**) Percentage of nuclei in each cluster determined by group. (**D**) Violin plots showing normalized expression levels of select differentially expressed genes in myonuclei populations. (**E**) Violin plots showing z-score transformed expression level of select marker genes used to determine cluster identity. (**F**) Gene set enrichment analysis (GSEA) of up-and down-regulated genes in PAD nuclei populations. (**G**) Representative images and quantification of the myosin heavy chain fiber type distributions from non-PAD and PAD (n=12 non-PAD, n=15 PAD). Statistical analysis in Panel G was performed using an unpaired two-tailed Student’s *t*-test when data if data were normally distributed or with a Kolmogorov-Smirnov test if found to have non-normal distribution.

**Table 1:** Physical and Clinical CharacterisGcs of Human PaGents.

| Characteristic | Non-PAD<br>(N=12) | PAD<br>(N=20) | P value<br>(X<sup>2</sup> or t-<br>test) |
| --- | --- | --- | --- |
| Mean age (SD) – yr. | 71.4 (9.2) | 67.8 (7.4) | 0.236<sup>A</sup> |
| Female sex – no. (%) | 5 (42) | 7 (35) | 0.999 |
| Body Mass Index (BMI, kg/m<sup>2</sup>) – (SD) | 29.7 (2.8) | 26.9 (5.9) | 0.136 |
| Ankle-brachial index (ABI) – (SD) | 1.08 (0.09) | 0.62 (0.15) | **5.02E-10** |
| Medical history – no. (%) |  |  |  |
| Diabetes mellitus type I or II | 4 (33) | 10 (50) | 0.584 |
| Hypertension | 6 (50) | 17 (85) | 0.084 |
| Hyperlipidemia | 5 (42) | 17 (85) | **0.030** |
| Coronary artery disease | 0 (0) | 11 (55) | **0.005** |
| Congestive Heart Failure | 0 (0) | 2 (10) | 0.708 |
| Tobacco use – no. (%) |  |  |  |
| Former smoker | 4 (33) | 14 (70) | 0.097 |
| Current smoker | 0 (0) | 3 (15) | 0.434 |
| Medication used – no. (%) |  |  |  |
| Aspirin | 4 (33) | 6 (30) | 0.841 |
| Statin | 5 (42) | 15 (75) | 0.131 |
| ACE inhibitor | 3 (25) | 5 (25) | 0.671 |
| Anticoagulant | 2 (17) | 10 (50) | 0.131 |
| Antiplatelet | 1 (8) | 9 (45) | 0.076 |
| Cilostazol | 0 (0) | 5 (25) | 0.167 |
| X<sup>2</sup> analysis was performed to determine differences in population proportions. SD, standard deviation. |  |  |  |

To compare non-PAD and PAD muscles, dataset integration was performed, resulting in a total of 57,364 nuclei for bioinformatic analysis and uniform manifold approximation and projection (UMAP) was used to visualize and resolve different nuclear populations (**Figure 1B**). We identified 12 unsupervised clusters of nuclei based upon their transcriptional profiles and assigned identioes by examining normalized expression of values of top markers and known marker genes for each cluster (**Figure 1C, E**) and the relative percentages of each nuclei population are shown in **Figure 1C**. We identified three different myonuclei populations using established marker genes resulting in clear detection of known myonuclei classes such as Type I (*MYH7*^+^, *TNNT1*^+^, *ATP2A2*^+^), Type II (*MYH1*^+^, *MYH2*^+^*, ATP2A1^+^, TNNT3*^+^, *MYL1*^+^), and Hybrid I-II myonuclei that expressed markers of both fast (Type II) and slow (Type I) myonuclei. Other identified clusters included Regenerating myonuclei (*MYH3^+^, TNNT3*^+^, *RUN×1*^+^), MuSCs (*PAX7*^+^), Neuromuscular Junction (NMJ)-associated myonuclei (*CHRNA1*^+^, *CHRNG^+^*, *MUSK*^+^), Neural cells (*SCN7A^+^, PAK3^+^*), Endothelial cells (*PECAM1*^+^), Fibro-adipogenic progenitors cells (FAPS) (*PDGFRA*^+^, and *GLI3*^+^), Smooth muscle cells (*MYH11*^+^, *ACTA2*^+^, *CARMN*^+^), Macrophages (*CD163*^+^), and other Immune cells (*PTPRC*^+^) (**Figure 1E**). Additional feature plots of the expression levels of select genes are shown in **Supplemental Figure 1**. A list of marker genes for each cluster can be found in **Supplemental Dataset 1**.

Quantification of the relative abundance of nuclei populations identified several expected differences including PAD muscles having a lower percentage of Type I myonuclei, MuSCs, and Endothelial cells, but a high proportion of Type II and Regenerating myonuclei, as well as NMJ-associated myonuclei when compared to non-PAD muscles (**Figure 1C**). Differentially expressed gene analysis performed on Type I, Type II, and Hybrid myonuclei uncovered upregulation of *ANKRD1*, *PPP1R27, BAG3, FOXO1,* and *HSPB8,* genes involved in stress response, atrophy, and autophagy,^44, 45^ as unique signatures to PAD muscle **(Figure 1D, Supplemental Figure 2)**. Moreover, downregulated genes in PAD myonuclei included *VEGFA*, *NOS1*, and *ESRRG* which collectively are known regulators of angiogenesis. Gene set enrichment analysis of PAD myonuclei unveiled consistent pathways enriched with upregulate genes related to ‘*cellular response to external stimulus’*, ‘*cellular response stress’* and ‘*regulation of autophagy*’ across all three myonuclei populations **(Figure 1F).** Additionally, these populations were also enriched for genes involved in ‘*positive regulation of cell death’*, ‘*response nutrient levels’* and ‘*cellular responses to oxygen levels’* confirming that these myonuclei were derived from an ischemic microenvironment. Down-regulated GO terms in PAD myonuclei included ‘*blood circulation*’, ‘*muscle contraction*’, and ‘*cellular respiration*’ which are consistent with prior studies documenting muscle weakness^6, 46, 47^, mitochondrial impairment^22, 28, 30, 35, 48, 49^, and deficient vascular supply^50^ in the PAD condition.

Next, we perform immunohistochemistry analysis on muscle specimens from non-PAD participants and PAD patients. Representative images and quantification of muscle fiber types confirmed the observations of decreased Type I myofibers and increased Type IIa and Hybrid myofibers when staining for specific myosin heavy chain isoforms (**Figure 1G**). However, it is worth ntiong that the non-PAD muscles had few Hybrid myofibers according to immunohistochemical analysis, but ∼10% of the total nuclei were found to express mRNA’s associated with both Type I and II myonuclei. This observation indicates that myofibers contain additional regulation of myosin heavy chain expression at the protein level. Analysis of myofiber cross-sectional area (CSA) within each fiber type revealed significant atrophy in PAD patient muscle across all identified fiber types but was displayed the largest effect size in Type I and Type IIb myofibers (**Figure 1G**).

### PAD paGents display unique myonuclei populaGons and transcripGonal profiles associated with atrophy, metabolic dysfuncGon, and stress response

To evaluate whether PAD mature myonuclei populations exhibit unique transcriptional signatures, we next performed unsupervised clustering on Type I, Type II, and Hybrid I-II myonuclei populations independently. A total of 26,190 Type I myonuclei from both non-PAD and PAD integrated datasets were subclustered to reveal ten distinct Type I (slow) myonuclei populations **(Figure 2A)**. Within each subcluster of Type I myonuclei, we performed gene set enrichment analysis to uncover processes that are differentially enriched in non-PAD and PAD muscles (**Figure 2B**). Within these subclusters, common deficiencies in gene expression in PAD Type I myonuclei were related to oxidative metabolism, muscle contraction, and vascular processes. In contrast, common upregulated pathways were primarily related to stress response processes. For example, feature plots of Type I myonuclei demonstrate significantly elevated expression of *ANKRD1* (also known as *CARP*), a myofibrillar stress response gene associated with cardiac and skeletal myopathies^51, 52^, as well as the hypoxia sensitive gene *EGR1*, in PAD muscle compared to non-PAD muscle (**Figure 2C**). Differential gene expression analysis also revealed an upregulation of *TRIM63* in PAD Type I myonuclei, also known as MuRF1, which is a master regulator of muscle mass ^53^ and known ubiquitin ligase involved in muscle atrophy (**Supplemental Figure 3A,B**). Of these subclusters, Clusters 7 was enriched in PAD muscle specimens, whereas Cluster 10 was predominantly derived from non-PAD muscle. Gene set enrichment analysis of Cluster 7 indicated an upregulation of genes associated with ‘*regulation of DNA repair*’ and downregulation genes involved in ‘*cytosolic calcium ion transport*’, ‘*vesicle-mediated transport in synapse’,* and ‘*vascular process in circulatory system’* likely indicating these Type I myonuclei were derived from areas of the slow myofiber undergoing denervation (**Figure 2B**). Cluster 7 also displayed significantly lower levels of expression of *MCU* and *VEGFA* compared to non-PAD (**Supplemental Dataset 2**), consistent with the GSEA results. Cluster 10, which was largely absent in PAD muscle, was characterized by high expression of Collagen Q (*COLQ*), a collagen species that anchors the acetylcholinesterase to the basal lamina in the neuromuscular junction via interactions with the MuSK receptor^54^. Additional genes which displayed elevated expression in PAD Type I myonuclei included the stress-inducible chaperone *HSPB1,* the orphaned nuclear receptor *NR4A1*, and the KLF transcription factor 13 (*KLF13*) (**Supplemental Figure 2B**).

**Figure 2.**
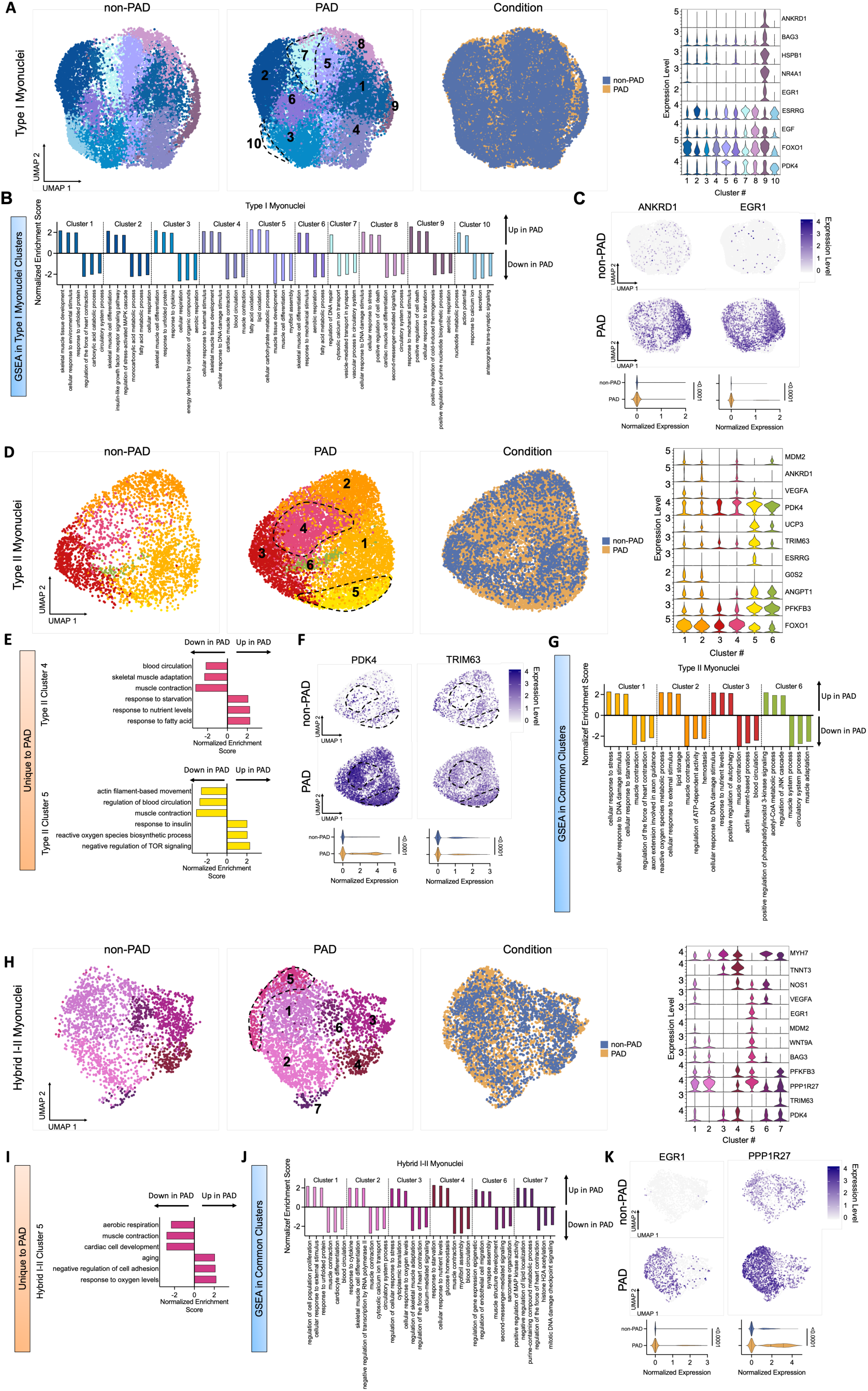
Subclustering analysis of myonuclei populaGons reveals transcripGonal signatures and myonuclei populaGons that are unique to PAD paGents. (A) UMAP visualizations of Type I myonuclei in non-PAD controls, PAD patients, and integrated UMAP by condition, as well as violin plots showing normalized expression levels of select genes. (B) GSEA terms of up-and down-regulated genes in Type I myonuclei subclusters. (C) Feature plots of *ANKRD1* and *EGR1* upregulated expression in Type I PAD myonuclei. (D) UMAP visualizations of Type II myonuclei in non-PAD controls, PAD patients, and integrated UMAP by condition, as well as violin plots showing normalized expression levels of select genes. (E) GSEA terms of clusters four and five Type II myonuclei. (F) Feature plots of significantly upregulated genes in PAD, *PDK4* and *TRIM63*, within Type II myonuclei. (G) GSEA terms of up-and down-regulated genes in clusters of Type II myonuclei found in both PAD and non-PAD muscle. (H) UMAP visualizations of Hybrid I-II myonuclei in non-PAD controls, PAD patients, and integrated UMAP by condition, as well as violin plots showing normalized expression levels of select genes. (I) GSEA terms in cluster five of Hybrid I-II myonuclei. (J) GSEA terms of common clusters of Hybrid I-II myonuclei. (K) Feature plots of select significantly upregulated genes in PAD (*EGR1* and *PPP1R27*) Hybrid I-II myonuclei.

A total of 11,943 Type II myonuclei were subclustered, resulting in six nuclei populations **(Figure 2D)**. Of these six subclusters, we found Clusters 4 and 5 to be predominantly present in PAD muscle (**Figure 2D**). Gene set enrichment analysis of these PAD-specific subclusters revealed enrichment of genes involved in ‘*response to starvaMo*n’ in Cluster 4 and ‘*negative regulation of TOR signaling*’ in Cluster 5, whereas both clusters exhibited deficiencies related to ‘*blood circulation*’ (**Figure 2E**). Consistent with these terms, both clusters displayed a significantly higher level of expression of *PDK4* and *TRIM63* in PAD muscles (**Figure 2F**), as well as *FOXO1* **(Supplemental Figure 3C,D)** genes all mechanistically linked to muscle atrophy^53, 55^. Cluster 4 from PAD patients was also enriched for *VEGFA* (**Supplemental Figure 3E,F**), a known regulator for angiogenesis in muscle. In addition to exploring PAD-specific subclusters, we also performed gene set enrichment analysis on Type II myonuclei clusters that were observed in both non-PAD and PAD muscles (**Figure 2G, Supplemental Dataset 3**). Globally, PAD Type II myonuclei displayed higher expression of uncoupling protein 3 (*UCP3,* **Supplemental Figure 2C,D**) which may contribute to the impaired oxidative metabolism commonly reported in PAD patients^22, 28, 30, 35, 56, 57^. Relatedly, several clusters in PAD muscle were enriched *for MDM2* (**Supplemental Figure 2C,D**), a negative regulator of p53 which has also been shown to negatively regulate mitochondrial respiration independent of p53 via transcriptional regulation of NADH-dehydrogenase 6 (*MT-ND6*)^58^. Intriguingly, deletion of *MDM2* in the skeletal muscle of mice has been shown to enhance muscle mitochondrial respiration and treadmill running performance in mice exposed to mild hypoxia^58^. Subcluster 5 was also found to be enriched with *ESRRG* expression (**Supplemental Figure 3C,D**), which has been shown to enhance ischemic angiogenesis and oxidative function of muscle in murine PAD models^59, 60^. Coincidentally, PAD Type II myonuclei also displayed substantial upregulation of *PFKFB3* (**Supplemental Figure 2D**), a known stimulator of glycolysis which is also linked to ischemic myopathy^61^.

Unsupervised clustering of 5,494 Hybrid I-II myonuclei generated seven distinct Hybrid myonuclei populations **(Figure 2H)**. Of these subclusters, Cluster 5 was found to be derived predominantly from PAD muscle specimens. Gene set enrichment analysis of Cluster 5 indicated an upregulation of genes involved with the ‘*response to oxygen levels*’, ‘*negative regulation of cell adhesion’,* and ‘*aging*, whereas the primary down regulated gene sets were related to ‘*muscle contraction*’ and ‘*aerobic respiration*’ (**Figure 2I**). Gene set enrichment analysis on Hybrid myonuclei clusters that were observed in both non-PAD and PAD muscles is shown in **Figure 2J**. Interestingly, clusters 1, 2, 5, and 7 in PAD muscle were enriched for *EGR1* and *PPP1R27* when compared to non-PAD (**Figure 2K**), suggesting these sub-populations of hybrid myonuclei may have been derived from hypoxia/stress-responsive regions of the muscle specimen. PAD hybrid myonuclei as displayed aberrant expression of the autophagy-related gene *BAG3*^62–66^ (**Supplemental Figure 2E,F)**, which is known to promote chaperone-mediated autophagy and maintenance of sarcomere structure ^67^, however, mutations in *BAG3* have been mechanisocally linked to ischemic limb myopathy in mice^68^. Datasets containing top marker genes, DEGs, and gene set enrichment analysis results for myonuclei subclusters can be found in **Supplemental Dataset 2** and **Supplemental Dataset 3** respectively.

### Increased transcriptome diversity and altered trajectory of MuSCs and regeneraGng myonuclei in PAD

Next, we examined explored whether PAD patient muscles had differences in the muscle stem cell (MuSC; also known as satellite cell) and regenerating myonuclei populations. A total of 1,906 MuSCs were present (909 nonPAD, 997 PAD) in our dataset. Differential gene expression analysis of the MuSC’s revealed a total of 145 differentially expressed genes (adjusted *P*-value < 0.05 and Log2FC > 0.25) between PAD and non-PAD. Of those, 102 were downregulated in PAD and 43 upregulated **(Figure 3A, Supplemental Dataset 1)**. Gene set enrichment analysis indicated that PAD MuSC’s were enriched for genes involved in *‘response to xenobiotic stimulus’*, *‘regulation of cell population and proliferation’, ‘response to oxidative stress’,* and *‘negative regulation of apoptotic process’* (**Figure 3B**). There were 3,578 regenerating myonuclei (1,063 from non-PAD and 2,515 from PAD muscle) indicating a greater relative abundance (5% vs.7% respectively) of regenerating nuclei in PAD patient muscle. DEG analysis of the regenerating myonuclei revealed a total of 1108 differentially expressed genes (adjusted *P*-value < 0.05 and Log2FC > 0.25) between PAD and non-PAD. Of those, 523 are downregulated in PAD and 585 upregulated in PAD muscle **(Figure 3C, Supplemental Dataset 1)**. Gene set enrichment analysis indicated that PAD regenerating myonuclei were enriched for genes involved in *‘response to growth factor’, ‘positive regulation of cell differentiation’, and ‘cellular response to oxygen levels’* (**Figure 3D**). Next, we performed unsupervised subclustering of the MuSC and regenerating myonuclei populations in non-PAD and PAD muscles. In non-PAD muscles, two clusters of MuSC’s were identified with both established satellite cell markers (*PAX7*, *MEG3*, and *MEG8*) and four clusters of regenerating myonuclei enriched with *MEF2C* (**Figure 3E**). In contrast, we found three clusters of MuSC’s and six clusters of regenerating myonuclei in PAD muscle indicating a greater diversity in transcriptional profiles is present in these cell populations (**Figure 3E**). Interestingly, PAD MuSC’s displayed significantly higher expression of several genes associated with the activator protein 1 (AP-1) transcription factor including *FOS*, *FOSB*, *JUN*, and *JUNB* **(Figure 3A,E,F**). Feature plots of *FOS* expression revealed a unique subpopulations of PAD MuSC’s and Regenerating myonuclei that were enriched for *FOS* (**Figure 3F**). Elegant work from a previous study involving non-ischemic muscle injury models demonstrated that this *FOS*^+^ population likely represents an activated/pro-regenerative MuSC’s and that deletion of *FOS* in satellite cells impaired muscle regeneration^69^. Nictionamide N-methyltransferase (*NNMT)*, which plays a role in cellular metabolic and epigenetic regulation, was also significantly upregulated in PAD MuSC’s and Regenerating myonuclei and displayed cluster specific enrichment in both cell types (**Figure 3F**). Notably, pharmacological inhibition of *NNMT* in skeletal muscle has been shown to enhance regeneration and improve MuSC function^70^. Of further interest, nictionamide, the substrate for NNMT, also serves as a precursor to the NAD^+^ salvage pathway which has been targeted as a potential therapeutic in a clinical trial for PAD patients involving nictionamide riboside treatment (NCT03743636). Other genes that were found to display unique expression patterns in PAD MuSC’s and Regenerating myonuclei were *FKBP5*, *DUSP1*, *ANKRD1*, and *MDM2* (**Figure 3F**). Interestingly, *FKBP5* is a peptidyl proyl isomerase that has been linked mechanistically to myogenesis^71^. *DUSP1*, a dual-specificity protein phosphatase involved in regulation of the cell cycle and apoptosis, was also upregulated in PAD and has been associated with atrophy due to impaired myoblast differentiation in cancer cachexia^72^. *ANKRD1* was also uniquely expressed in two subpopulations of Regenerating myonuclei in PAD muscle (**Figure 3F**). Previous studies have shown that *ANKRD1* expression is upregulated in response to denervation and could be involved in pathological remodeling of myofilaments ^52, 73, 74^. Finally, MDM2, a ubiquitin ligase known to degrade p53 ^75^, was also upregulated in PAD Regenerating myonuclei (**Figure 3F**). Notably, MDM2 has also been reported to be imported into mitochondria and negatively regulate respiratory function ^58^.

**Figure 3.**
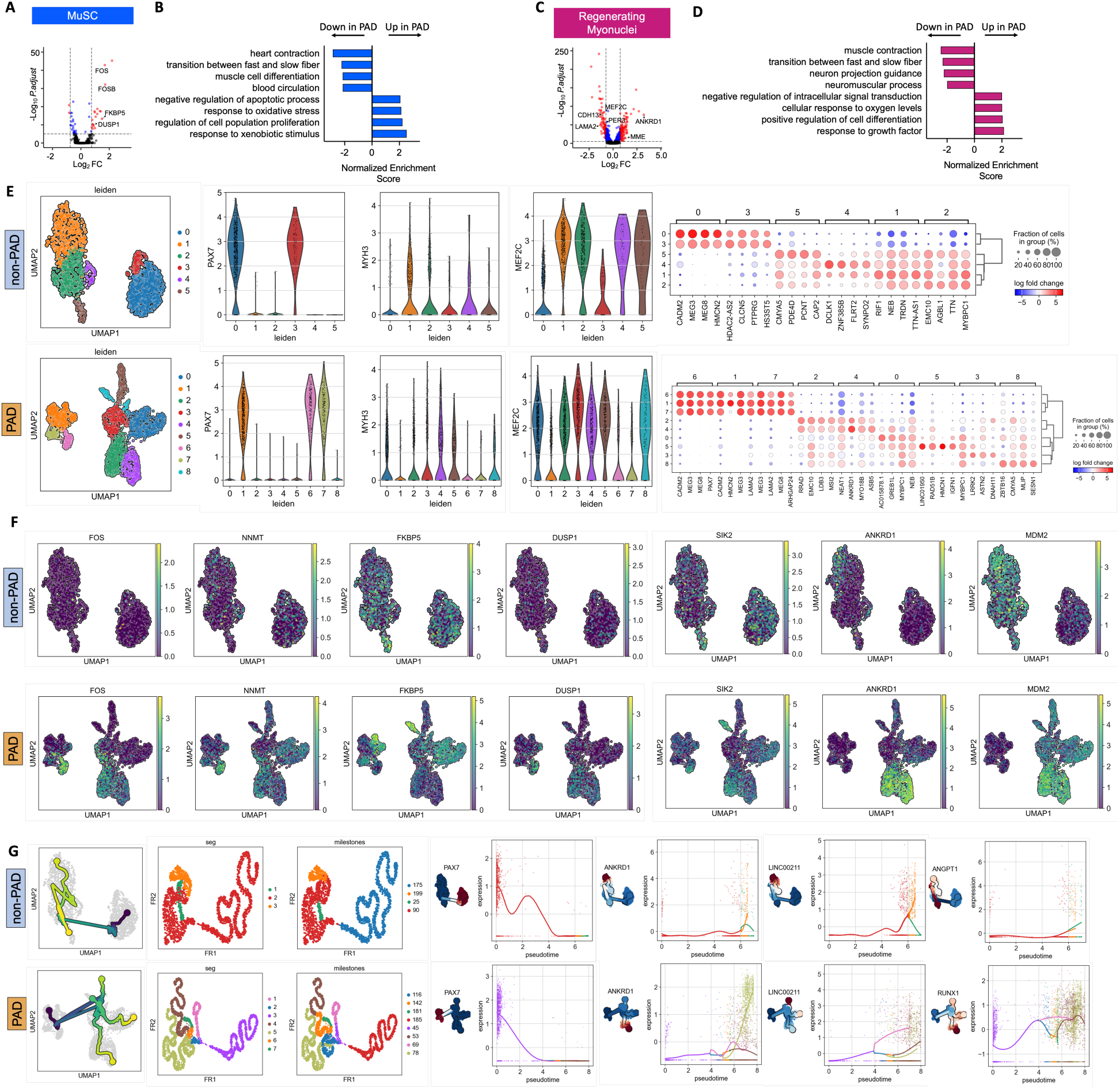
Increased transcripGonal diversity and altered trajectory of PAD MuSCs and regeneraGng myonuclei. (**A**) Volcano plot of significantly up-and down-regulated genes in PAD MuSC nuclei. (**B**) Significant GSEA terms up-and down-regulated in PAD MuSC nuclei. (**C**) Volcano plot of up-and down-regulated genes in PAD regenerating myonuclei. (**D**) Significant GSEA terms up-and down-regulated in PAD regenerating myonuclei. (**E**) UMAP visualizations of non-PAD and PAD MuSC and regenerating myonuclei with violin plots of top markers genes for subclusters. Dot plots of the top four non-PAD and PAD cluster markers with greatest log2 fold change. (**F**) Feature plots showing expression of select genes in non-PAD and PAD. (**G**) Trajectory plots overlayed on UMAPs for PAD and non-PAD MuSC and regenerating myonuclei. Visualization of tree analysis showing identified significant ‘segments’ and ‘milestones’, as well as branch-specific gene expression of select genes with changes across pseudtiome.

Next, we performed trajectory analysis on these cell populations and characterized genes that displayed significant expression changes in pseudtiome. For this analysis, the root was set using *PAX7* and trajectory analysis revealed three distinct segments in non-PAD and seven segments in PAD (**Figure 3G**). This resulted in PAD muscles having three branches of regenerating myonuclei, whereas non-PAD muscles appeared to have a single branch. Analysis of genes that displayed significant changes in expression across pseudtiome revealed 464 significant genes in non-PAD muscle and 1,522 Significant genes in PAD muscle (**Supplemental Dataset 4**). Examples of genes with significant changes in pseudtiome expression included *PAX7*, *ANKRD1*, *LINC00211*, *RUN×1*, and *ANGPT1* (**Figure 3G**). Future work is needed to determine if the genes with significant expression changes in pseudtiome serve functional roles in regulating myogenesis or muscle repair in PAD.

### Non-PAD and PAD endothelial and smooth muscle cells have similar transcripGonal profiles

To understand the transcriptional similarities and differences of non-PAD and PAD vascular cell populations, both smooth muscle and endothelial cells were subset out for each condition. Differential gene expression analysis for endothelial cells and smooth muscle cells are shown in **Supplemental Dataset 1**. Gene set enrichment analysis of endothelial cells revealed upregulation of genes related to ‘*inflammatory response’*, ‘*vasculature development’*, ‘*circadian regulation of gene expression*’ and ‘*response to cytokine’* in PAD patients. Expectedly, genes related to ‘*negative regulation of epithelial cell proliferation’, ‘circulatory system processes’,* and *‘blood circulation’* were downregulated in PAD **(Figure 4A, Supplemental Dataset 1).** We also perform immunohistochemistry to quantify the number of capillaries (PECAM1/CD31 staining) and arterioles (Myh11 staining) in non-PAD and PAD muscles. Despite the hypoxic condition of the PAD limb environment, PAD patients had similar capillary density (*P*=0.7267) but lower arteriole density (*P*=0.0196) compared to non-PAD participants (**Figure 4B**).

**Figure 4.**
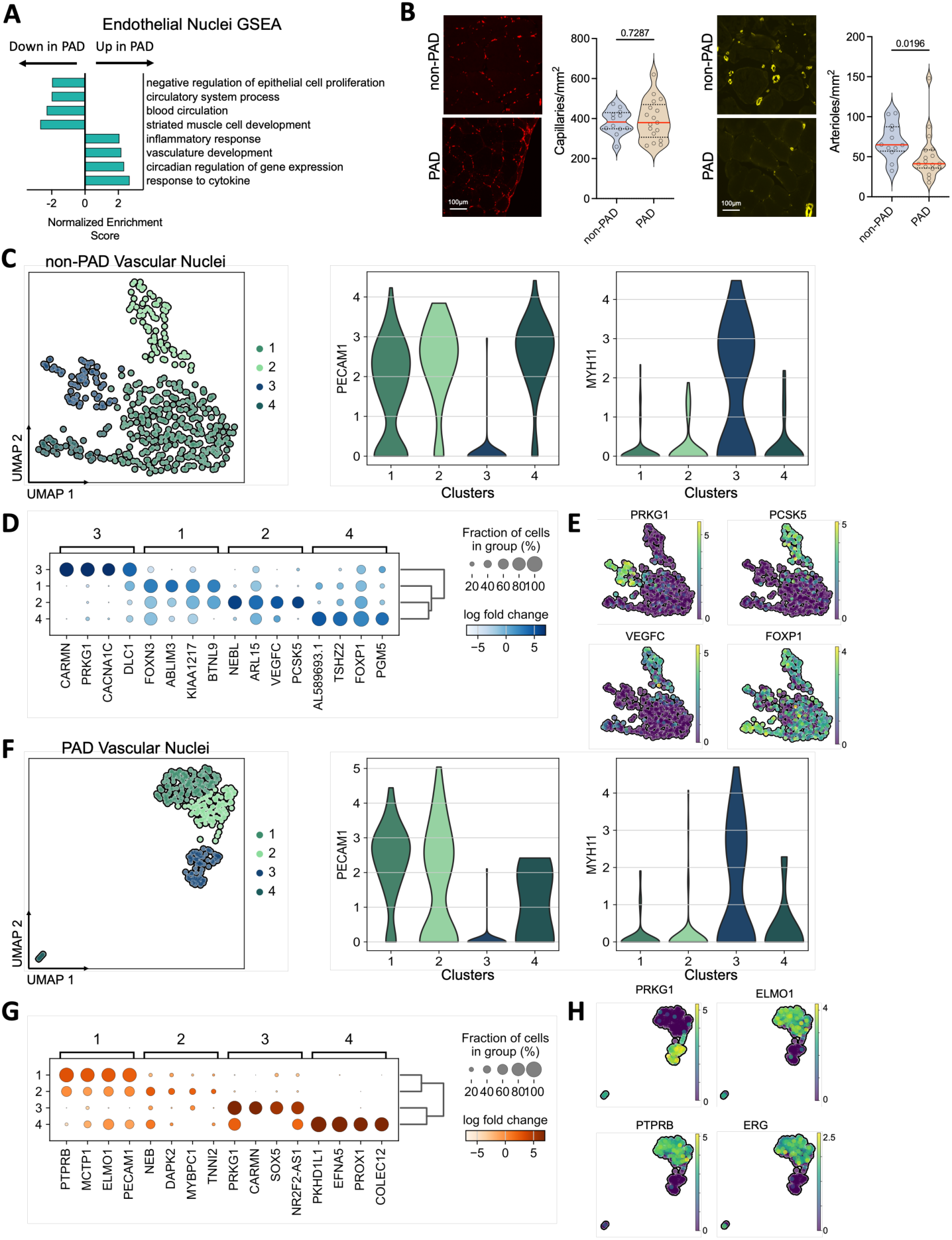
Analysis of endothelial and smooth muscle cell nuclei in non-PAD and PAD gastrocnemius muscle. (**A**) GSEA terms of up-and down-regulated genes in PAD endothelial cell nuclei. (**B**) Representa?ve images and quan?fica?on of capillary (PECAM1/CD31 staining) and arteriole (MYH11 staining) density in non-PAD and PAD muscle (n=12 non-PAD, n=17 PAD). Capillary density was analyzed using an unpaired two-tailed Student’s *t*-test, while arteriole density was analyzed using a -Smirnov test. (**C**) UMAP visualiza?on of non-PAD vascular nuclei and violin plots of known marker genes *PECAM1* and *MYH11*. (**D**) Dot plot of the top four cluster markers in non-PAD vascular nuclei. (**E**) Feature plot visualiza?on of select marker genes expression in non-PAD vascular nuclei. (**F**) UMAP visualiza?on of PAD vascular nuclei and violin plots of known marker genes *PECAM1* and *MYH11*. (**G**) Dot plot of the top four marker genes in PAD vascular nuclei. (**H**) Feature plot visualiza?on of select marker genes expression upregulated in PAD vascular nuclei.

To further characterize the vascular cell populations, cells were subclustered to identify distinct populations present in each condition. A total of 461 non-PAD vascular nuclei revealed four potential subpopulations. We classified cluster three as smooth muscle (*MYH11^+^*) and the others were clearly endothelial cells (*PECAM1^+^*) **(Figure 4C)**. To delineate each cluster further, the top expressed genes were calculated **(Supplemental Dataset 5)**. The top four genes for each cluster were plotted and tree lineage analysis revealed greatest transcriptional similarity between clusters two and four **(Figure 4D).** High expression of *CARMN* and *PRKG1* further validated the classification of smooth muscle cells in cluster three^76, 77^ **(Figure 4D,E).** Proprotein convertase 5/6, *PCSK5,* was uniquely expressed in cluster two of non-PAD endothelial cells. Inactivation of *PCSK5* in endothelial cells has been shown to result in cardiovascular hypertrophy associated with reduced collagen production, IGF-1/Akt/mTOR signaling, increased vascular stiffness, and enhanced autophagic activation^78^. Additionally, *VEGFC*, arose as one of the top markers for cluster two non-PAD endothelial cells **(Figure 4D,E)**. Uniquely, *FOXP1*, a gene that has been identified to stimulate angiogenesis following myocardial infraction ^79^, showed highest expression in cluster four but was also expressed in the other three vascular clusters **(Figure 4D,E)**. PAD patient muscle contained 308 high quality vascular nuclei that resulted in four distinct clusters **(Figure 4F)**. Similar to non-PAD, cluster three was characterized as smooth muscle cells (*MYH11^+^*), while clusters one, two, and four were endothelial cells (*PECAM1^+^*) **(Figure 4F)**. Gene expression analysis of each cluster and tree lineage revealed strongest transcriptional similarities between clusters one and two **(Figure 4G)**. Like non-PAD, smooth muscle cells had high expression of *CARMN* and *PRKG1*, validating this cluster as smooth muscle cells in PAD **(Figure 4G,H)**. Engulfment and cell mtiolity 1 (*ELMO1)*, which showed high expression in cluster one, has been implicated in angiogenesis and stabilization of the endothelium^80^, **(Figure 4G,H)**. *PTPRB*, a gene that encodes a transmembrane protein known to regulate blood vessel remodeling and angiogenesis^81^, was also a top expressed gene in endothelial cell clusters one and two from PAD muscle (**Figure 4H**). Interestingly, *PTPRB* knockdown in epithelial cells resulted in higher activation of FGFR, implicating *PTPRB* plays an important role in regulating branching morphogenesis through the FGF-FGFR signaling pathway^82^. Finally, expression of *ERG* (a member of the erythroblast transformation-specific family of transcription factors) was highly enriched in cluster one of PAD endothelial cells (**Figure 4H**). *ERG* has a critical for endothelial homeostasis, migration, and angiogenesis^83, 84^. Together, these findings suggest the PAD vascular populations are expressing, at the transcriptional level, genes important for mediating angiogenesis, however, immunohistochemistry analysis of non-PAD and PAD muscle suggests that these processes are not manifested as increased small vessel density within the affected skeletal muscle (**Figure 4B**).

### PAD muscle is characterized by increase fibrosis and altered transcripGonal signatures of fibro-adipogenic progenitor cells (FAPs)

FAPs are a mesenchymal stem cell population that reside within skeletal and have the ability to differentiate into fibroblasts or adipocytes, but are believed to lack the ability to directly form myoblasts^85, 86^. Upon muscle injury, FAPs become activated, proliferate, and can promote MuSC-mediated regeneration via paracrine signaling events ^85, 87^. However, in pathological conditions, the aberrant proliferation/differentiation of FAPs have been linked to skeletal muscle fibrosis^88^. In this regard, Masson’s Trichrome staining of PAD and non-PAD gastrocnemius muscles demonstrated that PAD patient muscles have significantly higher areas of fibrosis (*P*=0.0279, **Figure 5A**), confirming previous reports in the literature ^89–91^. Differential gene expression analysis of FAPs revealed 192 genes upregulated in PAD and 368 downregulated genes (adjusted *P*-value < 0.05 and Log2FC > 0.25) **(Figure 5B).** Collagen type IV alpha 1 (*COL4A1*) and alpha 2 (*COL4A2*) chains were upregulated in PAD FAPs nuclei (**Figure 5B**, Supplemental Dataset 1). Collagen type IV is a major component of the basement membrane in the extracellular matrix (ECM). In the context of skeletal muscle fibrosis, collagen is upregulated and takes up a large portion of muscle volume^92^. The upregulation of *COL4A1* and *COL4A2* may be contributing to excess production and deposition of collagen reported in PAD muscle^89^. Interestingly, there was significant downregulation of *EPHA7,* a gene that has been implicated in skeletal muscle regeneration^93^. Additionally, there is downregulation of *ITGA11* in PAD FAPs, a gene that is regulated by hedgehog signaling^94^ which has been linked to determining the cell fate of FAPs^95^. Gene set enrichment analysis of genes upregulated in PAD FAPs nuclei revealed terms related to *‘cell population proliferation*’, ‘*positive regulation of transcription by RNA polymerase II*’, ‘*response to cytokine*’, and interestingly, ‘*vasculature development’*. Downregulated terms in FAPs from PAD muscle included ‘*regulation of transforming growth factor beta receptor signaling pathway’*, ‘*neuron projection guidance*’, ‘*negative chemotaxis*’ and ‘*skeletal muscle adaptation*’ (**Figure 5C**).

**Figure 5.**
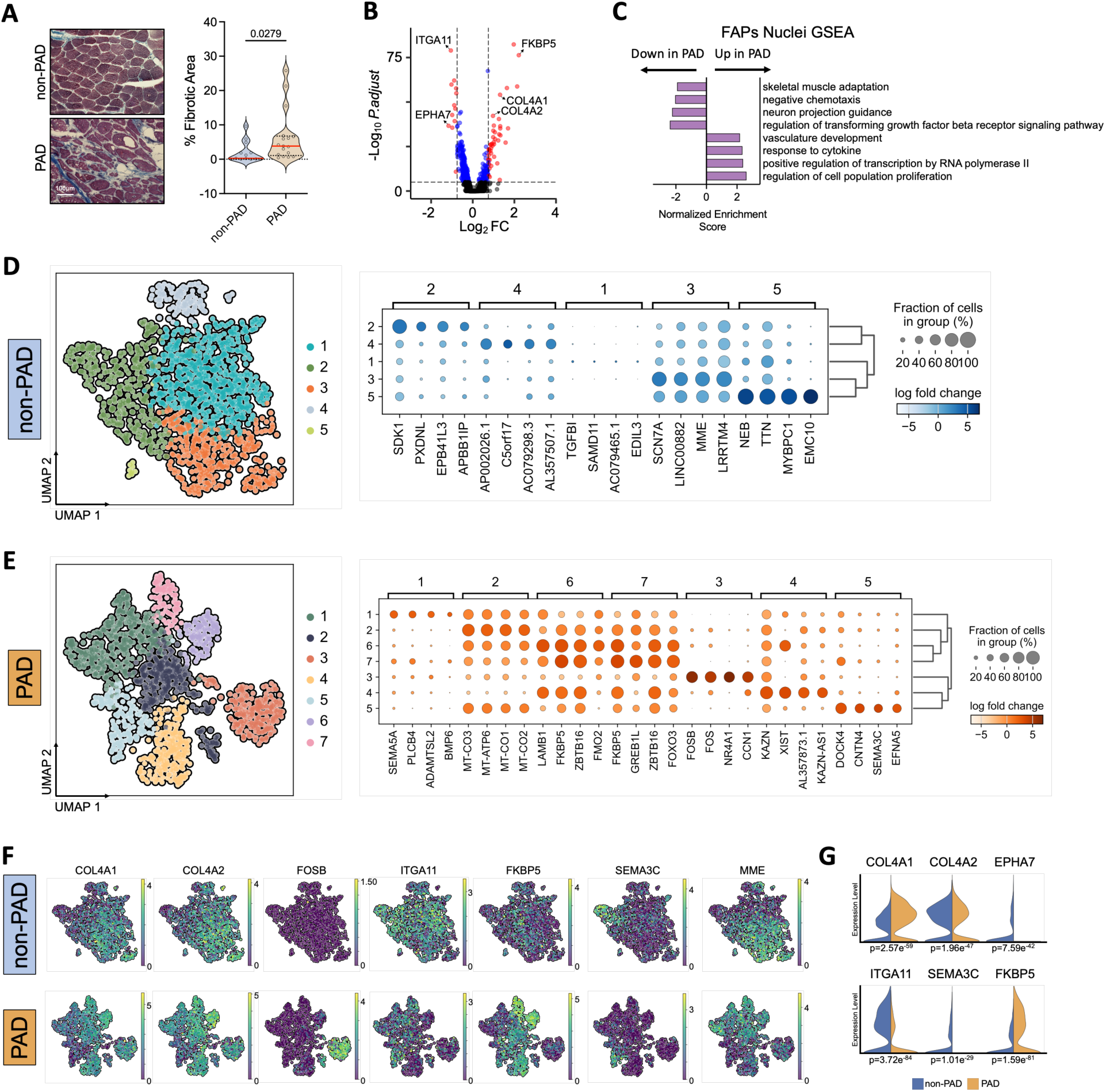
FAPs have disGnct transcripGonal signatures in PAD muscle. (**A**) Representative Masson’s trichrome images and quantification of fibrtioc area from non-PAD and PAD muscles (n=12 non-PAD, n=17 PAD) analyzed using a Kolmogorov-Smirnov test. (**B**) Volcano plot of significantly up-and down-regulated genes in PAD FAPs. (**C**) GSEA terms of significantly up-and down-regulated genes in PAD FAPs. (**D**) UMAP visualization and dot plot of the top four cluster markers in non-PAD FAPs. (**E**) UMAP visualization and dot plot of the top four cluster markers in PAD FAPs. (**F**) Feature plots of select DEGs in non-PAD and PAD FAPs. (**G**) Split violin plots of top DEGs presented with adjusted *P*-values.

To delineate how FAPs are impacted by the ischemic microenvironment in PAD, we subset the 2,880 non-PAD and PAD FAPs (*PDGFRA*^+^ nuclei) and independently subclustered each condition. In non-PAD participants, unsupervised clustering of 1,397 FAPs resulted in five distinct FAP populations, although cluster 5 was small in number (**Figure 5D**). Top marker genes for non-PAD FAP subclusters are shown in **Figure 5D**. In contrast, employing identical clustering parameters, 1,683 FAPs from PAD muscle produced seven transcriptionally distinct populations (**Figure 5E**). Some notable differences in expression in PAD FAPs were uncovered. For example, FOSB was highly expressed only in cluster 3 of PAD FAPs, but largely absent in non-PAD FAPs (**Figure 5F**). FKBP5 was also found to be enriched in four specific clusters of PAD FAPs (clusters 2, 4, 6, and 7), but displayed significantly lower expression in non-PAD FAPs (**Figure 5F,G**). In contrast, MME, a recently identified marker of adipogenic FAPs^96^, was expressed in both non-PAD and PAD FAPs but displayed cluster-specific enrichment in both conditions (**Figure 5F**). These findings demonstrate that the PAD patients exhibit greater transcriptional diversity in FAPs characterized by downregulation of genes that play a role in skeletal muscle regeneration and upregulation of genes related to fibrosis and remodeling of the extracellular matrix.

### PAD paGents exhibit alteraGons in inferred cellular communicaGon

Next, we explored ligand-receptor interactions using CellChat^43^ to determine how the PAD condition alters communication between the 13 different identified nuclei populations. A total of 1,486 ligand-receptor interactions were identified in non-PAD, whereas 1,208 interactions were detected in PAD muscle (**Figure 6A,B and Supplemental Figure 4A**). Overall, the inferred strength of interactions was greater in non-PAD compared to PAD, suggesting that the PAD condition disrupts cellular communication within muscle **(Supplemental Figure 4A)**. Circle plots depicting the global intercellular communication are shown in **Figure 6A**, where the line width represents the communication strength. A complete list of intercellular communication in non-PAD and PAD datasets can be found in **Supplemental Dataset 6**. Quantification of differential signaling interactions between non-PAD and PAD revealed an increase in the number of interactions between regenerating myonuclei and FAPs in PAD, whereas macrophages, endothelial, smooth muscle, and neural cells had decreased level of intercellular communication in PAD (**Figure 6B).** Interestingly, comparison of the major senders and receivers, FAPs were identified as the major outgoing source in both non-PAD and PAD, whereas the endothelial cell population received the most signals in both non-PAD and PAD (**Figure 6C**). In general, PAD muscles displayed a reduction in both outgoing and incoming interaction strength compared to non-PAD across nearly all nuclei populations suggesting that the ischemic microenvironment disrupts normal intercellular communication.

**Figure 6.**
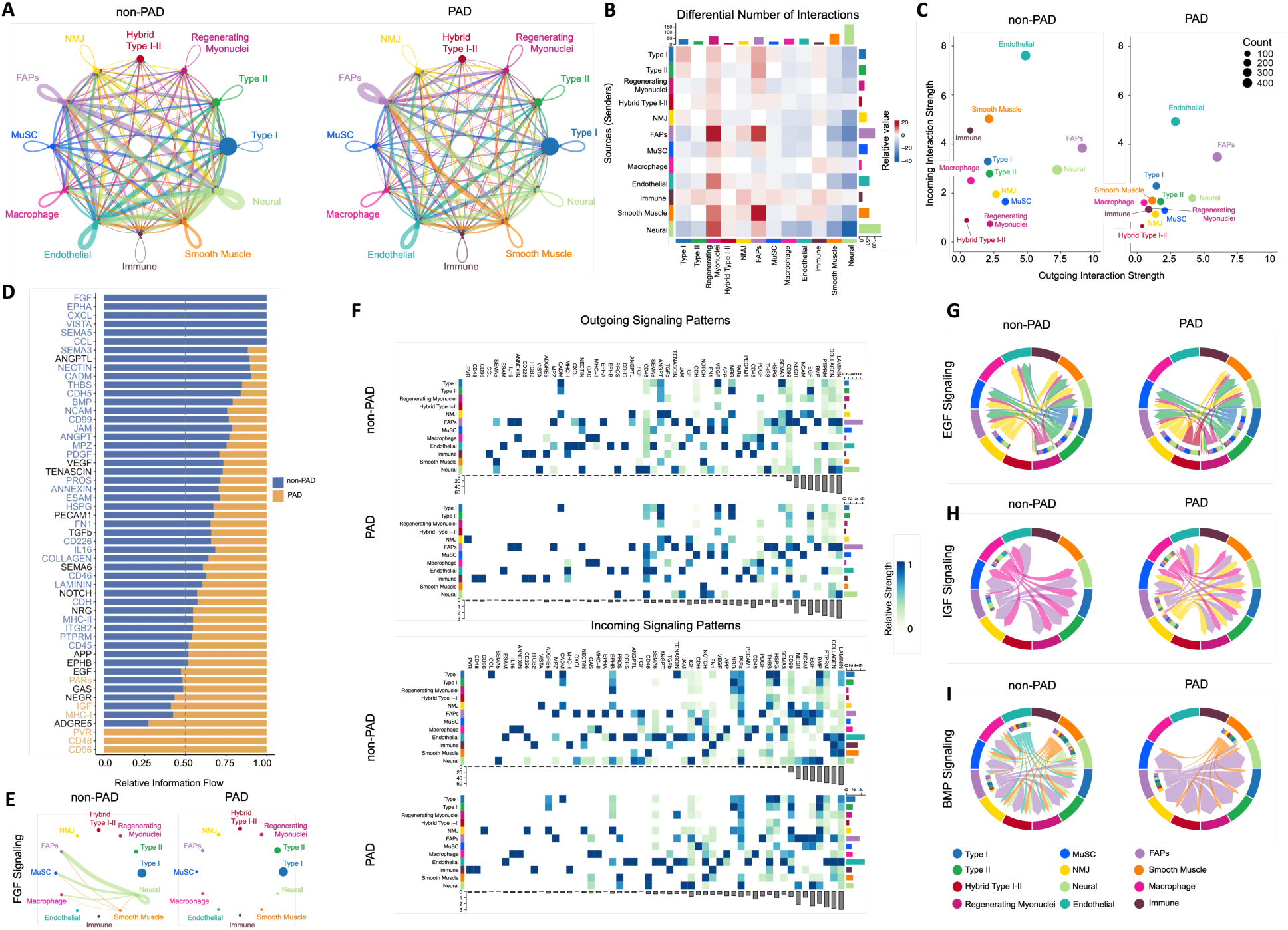
CellChat analysis predicts changes in intercellular communicaGon of PAD paGents. **(A)** Circle plots showing the overall intercellular communication occurring in non-PAD and PAD. Circle sizes represent the number of cells and edge width represents communication probability. (**B**) Differential number of interactions between non-PAD and PAD. Red represents increased number of signaling interactions in PAD compared to non-PAD. (**C**) Comparison of outgoing and incoming interaction strengths for all cells type in non-PAD and PAD. (**D**) Ranked significant ligand-receptor communications for relative information flow between non-PAD and PAD muscles. (**E**) Circle plots for FGF signaling communication in non-PAD and PAD. (**F**) Enriched outgoing and incoming signaling patterns according to cell type and signaling strength. (**G**) Chord plots for EGF signaling communication in non-PAD and PAD. (**H**) Chord plots for IGF signaling communication in non-PAD and PAD. (**I**) Chord plots for BMP signaling communication in non-PAD and PAD.

Significant signaling pathways between non-PAD and PAD were identified according to variations in information flow within inferred signaling networks **(Figure 6D)**. Enriched signaling pathways in non-PAD that were absent in PAD included FGF, EPHA and CXCL. Ligands *FGF1* and *CXCL12* were enriched in non-PAD **(Supplemental Figure 4)**. The potential importance of this observation is underscored by the known roles that fibroblast growth factor (FGF) signaling plays in promtiong angiogenesis, myogenesis and tissue regeneration in skeletal muscle. Non-PAD muscles expressed *FGF* ligands in both Smooth muscle and Neural cells where the primary interactions were with FAPs, MuSCs, and Macrophages (**Figure 6E**). However, FGF communication was absent in PAD **(Figure 6E).** Further comparative analysis of the outgoing and incoming signaling patterns between non-PAD and PAD allowed identification of signaling pathways and ligand-receptor interactions that exhibited distinct and similar signaling patterns **(Figure 6F)**. In both conditions, LAMININ and COLLAGEN which are important components of the extracellular matrix, arose as the signaling pathways with the greatest number of intercellular interactions **(Figure 6F, Supplemental Figure 4)**. Surprisingly, TGF beta signaling was completely lost between FAPs and Regenerating myonuclei in PAD, but present in non-PAD **(Supplemental Figure 4)**. The presence of EPHA and CXCL signaling in non-PAD muscle was dominated between FAPs, endothelial, and neural cells, but was again absent in PAD **(Supplemental Figure 4).** Epidermal growth factor signaling (EGF-EGFR ligand-receptor interactions) were markedly upregulated in PAD myonuclei populations but interestingly downregulated between regenerating myonuclei and endothelial cells **(Figure 6G)**. Epidermal growth factor receptor (EGFR), a member of the ErbB family, is traditionally linked to angiogenesis. However, previous work as shown that EGF signaling is not required for skeletal muscle angiogenesis in murine hindlimb ischemia models^97^. More recently, inhibition of EGF signaling was reported to promote a slow-oxidative muscle phenotype including increased mitochondrial content^98^. Given that EGF-EGFR is dominated by myonuclei populations in PAD and was found to be absent in endothelial cells, this finding highlights to potential for EGFR inhibitors to improve muscle pathology in PAD. Insulin-like growth factor (IGF) signaling was also upregulated in PAD, with unique communication coming from the nuclei associated with the neuromuscular junction (NMJ) in PAD patients **(Figure 6H)**. BMP signaling was downregulated in PAD skeletal muscle, with a loss of communication from endothelial cells observed **(Figure 6I)**. Notably, decreased BMP signaling has been reported in skeletal muscle of mice with cancer cachexia and was mechanistically linked to dismantling of the NMJ and myofiber denervation^99^, both phenotypes that would presumably contribute to poor muscle function and mobility impairment in PAD.

## DISCUSSION

In recent years there has been an emergence of information documenting the skeletal muscle pathology in lower limbs in patients with PAD^21, 27, 28, 30, 35, 57,100–105^, however the mechanisms driving lower limb functional impairment and mobility loss remain poorly understood. A few studies have employed wide-ranging ‘omics’ technologies, such as RNA sequencing and metabolomics, to uncover novel genes/pathways associated with PAD outcomes^106–108^, however these studies utilized blood specimens rather than muscle biopsies obtained from below the atherosclertioc lesion. Only three studies have leveraged unbiased RNA sequencing analysis on muscle specimens from PAD^28, 105, 109^. Results from these studies have consistently reported that muscle from PAD patients displays reduced expression of mitochondrial associated genes and increased levels of genes involved in hypoxia, inflammation, and autophagy. While much has been learned about muscle pathology in PAD from these studies, they all employed sequencing on RNA samples isolated from a small specimen obtained from the gastrocnemius muscle where the final RNA pool consists of material derived from a diverse range of cell types. Recent advances in single cell sequencing technologies have highlighted that skeletal muscle contains a diversity of cell populations that contribute to the overall health and function of muscle^31, 32, 34^. In this study, we applied single nuclei RNA sequencing to gastrocnemius muscle specimens obtained from non-PAD volunteers and patients with PAD with the objective of characterizing differences in cell/nuclei populations, transcriptional signatures, and intercellular communication in the PAD patient.

While all expected major cell types were present in PAD muscles, PAD patients displayed a relative increase in the proportion of type II (fast/glycolytic) myonuclei and decrease in type I (slow/oxidative) myonuclei compared to non-PAD which was observed in both snRNAseq and with immunohistochemistry (**Figure 1**). This slow-to-fast fiber type transition is commonly observed in conditions that involve muscle atrophy^110^. In regard to the latter, PAD myofiber areas were significantly smaller across all fiber types herein (**Figure 1H**), an observation consistent with previous work^111^. The observed fiber type distributions in the current study does not fully agree with a previous study that reported a trend toward increased proportions of hybrid Type I-II myonuclei in PAD, but similar relative proportions of Type I and Type IIa in PAD and non-PAD muscles^112^. Related to myofiber and muscle size, PAD muscles displayed significantly higher expression of known atrophy-promtiong genes (*FOXO1*, *GADD45A*, *TRIM63*)^113, 114^. Additionally, analysis of inferred intercellular communication revealed deficiencies in bone morphogenetic protein (BMP) signaling in myonuclei populations of PAD patients. Previous work has demonstrated that BMP signaling (via Smad1/5/8) is a fundamental signal that controls skeletal muscle mass^115^. Consequently, conditions that display muscle atrophy are characterized by deficient BMP signaling and rescue of BMP signaling has been shown to prevent/attenuate muscle atrophy^99^. Importantly, the study from White *et al*.^112^ reported a significant positive correlation between the Type I fiber area and walking performance in PAD, which confirms findings from Koutakis *et al.*^111^. At the whole muscle level, lower muscle area has also been associated with poor walking performance^25^. Taken together, these observations suggest that interventions that increase BMP signaling may have efficacy at increasing muscle/myofiber size that could confer improvements in mobility/walking performance in PAD.

In PAD myonuclei populations, common upregulated gene pathways included stress response, hypoxia, and autophagy which is consistent with work by Ferrucci and colleagues ^105^ using bulk RNA sequencing and proteomics analysis of PAD muscles. PAD myonuclei displayed high expression of *ANKRD1*, *BAG3*, and *HSPB8* (**Supplemental Figure 2**). *ANRKD1*, also known as muscle ankrin repeat protein 1 (*MARP1*), has been described as both a transcriptional factor^74^ and a stress-inducible myofibrillar protein that interacts with tion ^116^. Elevated expression of

*ANKRD1* has been previously documented in atrophying myofibers^52^ and failing heart/myocardium^51^ suggesting a potential mechanistic role in muscle dysfunction in PAD. *BAG3* and *HSPB8*, which have elevated expression in PAD, are both involved in chaperone-mediated autophagy and myofibrillar integrity^67, 68, 117, 118^. Interestingly, point mutations in *BAG3* have been mechanistically linked to ischemic muscle injury and recovery in murine PAD models^68^. Mutations in *BAG3* and *HSPB8* have been linked to hereditary skeletal myopathies^118, 119^. Unfortunately, the current study did not perform exome sequencing on PAD patients, so it was not possible to explore whether PAD patients exhibited mutations in any of these myopathy-associated genes.

A novel finding from this study is that PAD increases the transcriptional diversity of several cell/nuclei populations within skeletal muscle obtained from below the atherosclertioc lesion. For example, in both type II and hybrid I-II myonuclei populations, we identified subclusters that were only present in PAD muscle (Figure 2). These PAD-specific myonuclei populations displayed expression patterns related to atrophy, nutrient/oxygen deprivation, as well as alterations in mitochondrial and glycolytic metabolism. In addition to mature myonuclei, PAD muscle also displayed a greater diversity in the populations of MuSCs and regenerating myonuclei, an observation that likely represents the adaptive response to the repeated ischemia-reperfusion injuries that are suspected to occur in the affected limb muscle. Finally, the FAPs, a resident mesenchymal stem cell population, were also found to be more diverse in their transcriptome compared with FAPs from non-PAD muscle (Figure 5). In addition to displaying more transcriptional diversity, analysis of intercellular communication identified FAPs as a major signaling hub within human skeletal muscle. While the inferred communication was markedly different in FAPs from PAD muscle compared with non-PAD muscle, it should be noted that additional studies are necessary to establish functional impacts of altered FAPs communication in the PAD condition.

There are some limitations to this study that are worth noting. First, this was a cross-sectional analysis and the identified changes in gene expression cannot be causally linked to PAD patient outcomes. Moreover, the sample size was small and the clinical phenotype relatively homogenous, which precluded any type of sub-group comparisons that could help distinguish transcriptional differences across the wide range of symptomology in PAD. Herein, we performed RNA sequencing on single nuclei isolated from gastrocnemius muscle. This approach was chosen because of concerns about how enzymatic digestion and tissue dissociation procedures, which are required to obtain single cell suspensions from skeletal muscle, may impact gene expression. Furthermore, these digestion processes result in the loss of myofibers which are multinucleated and are not present in the final single cell suspensions. Nonetheless, there are some limitations to the single nuclei preparation employed herein. For example, myonuclei were the dominant population identified in all samples and there are cell populations within skeletal muscle that we could not identify that might play significant roles in PAD pathobiology (i.e., pericytes, tenocytes, Schwann cells, etc.). Additionally, transcripts identified in the represent those that were present within the nucleus. Thus, transcripts that are quickly exported to the cytoplasm could be under-represented in our dataset. Finally, sequencing depth is still a major limitation of both single cell and single nucleus RNA sequencing experiments and less abundant transcripts may not have been detected in our dataset but could still play important roles in muscle pathology in PAD. Despite these limitations, this analysis provides a detailed picture of the transcriptional differences associated with skeletal myopathy in PAD as a single nuclei resolution and may indicate new therapeutic targets/pathways that could be exploited to improve muscle function and mobility in PAD.

## CONCLUSIONS

Pharmacologic therapies for PAD remain limited and center on reducing the risk of cardiovascular events rather than improving limb function and mobility. Complicating therapeutic progress is a limited understanding of the mechanisms contributing to skeletal muscle pathology in PAD. This study employed snRNA sequencing to uncover novel nuclear populations and transcriptional differences that were unique to PAD muscle. This atlas gene expression in PAD provides a high-resolution understanding of the complex muscle pathology and aberrant intercellular signaling that is present in the ischemic gastrocnemius muscle.

### Funding Support

This study was supported by National Institutes of Health (NIH) grant R01-HL149704 (T.E.R.). K.K. was support by the American Heart Association grant POST903198. T.T. was support by National Institutes of Health (NIH) grant F31-DK128920.

### Author Disclosures

The authors have no conflicts, financial or otherwise, to report.

